# Single-Cell Transcriptomics reveals relaxed evolutionary constraint of spermatogenesis in two passerine birds as compared to mammals

**DOI:** 10.1101/2022.01.22.477241

**Authors:** J. Carolina Segami, Marie Semon, Catarina Cunha, Claudia Bergin, Carina F Mugal, Anna Qvarnström

## Abstract

Spermatogenesis is a complex process where spermatogonia develop into haploid, mobile sperm cells. The genes guiding this process are subject to an evolutionary trade-off between preserving basic functions of sperm while acquiring new traits ensuring advantages in competition over fertilization of female gametes. In species with XY sex chromosomes, the outcome of this trade-off is found to vary across the stages of spermatogenesis but remains unexplored for species with ZW sex chromosomes. Here we characterize avian spermatogenesis at single cell resolution from testis of collared and pied flycatchers. We find evidence for relaxed evolutionary constraint of genes expressed in spermatocyte cells going through meiosis. An overrepresentation of Z-linked differentially expressed genes between the two species at this stage suggests that this relaxed constraint is associated with the lack of sex-chromosome silencing during meiosis. We conclude that the high throughput of bird spermatogenesis, at least partly, is explained by relaxed developmental constraint.

## Introduction

Spermatogenesis, the biological process by which haploid sperm cells are created from diploid spermatogonia, is a highly complex process where at least seven somatic cell types and 26 morphologically distinct germ cell classes are involved^1^. Major chromatin remodeling occurs during spermatogenesis such as programmed double strand breaks, homologous recombination, chromosome synapsis, chromatin packing and chromosome inactivation^2^. As a result, the testis, where these processes occur, has the most complex transcriptome among all tissues including the brain^3,4^. Some of this complexity may be explained by the prevalent transcription occurring across large parts of the genome, which is thought to happen either as a none-adaptive side-effect of chromatin remodeling or as a mechanism of general “transcriptional scanning” that allow DNA repair^3,5,6^. However, the testis also harbors the highest number of tissue-specific expressed genes among all tissues in mammals, birds and insects^7–11^. A crucial question is to what extent genes with testis-specific expression reflect evolutionary innovations and species-specific differences in mating behaviors and sperm-egg interactions or are shared among divergent species and evolutionary constrained through effects of purifying natural selection to keep the complex process of spermatogenesis functional.

Analyses on the protein-coding sequences and transcription patterns show that testis-specific genes are among the most rapidly evolving genes^12,13^ and that de-novo genes often appear in the testis^13^. In addition, testis size^14,15^ and sperm morphology are astonishingly diverse among animal species^16^. This fast evolution is thought to be largely driven by sexual selection, mainly through sperm competition^17–19^ and cryptic female choice, the mechanisms by which females select the outcome of sperm competition^20–24^. Both these processes are post-copulatory events that occur when females have mated with multiple males, causing competition over fertilization success among ejaculates^19,25^.

The great diversity in spermatogenesis and sperm phenotypes across animal species may appear paradoxical in light of the central goal of this whole process being the same across species, i.e. to ensure the transmission of genetic material to the next generation. To achieve this goal, basic prerequisites always need to be fulfilled, including correct mitosis and meiosis and finally maturation of sperm cells that ensures their motility and ability to recognize and merge with the oocyte. Therefore the very basic processes of spermatogenesis are similar among different animals and there is evidence that the underlying genes are highly conserved^26,27^. Thus, spermatogenesis is subject to a combination of strong positive sexual selection through sperm competition and female cryptic choice as well as strong purifying natural selection to maintain central functions.

The balance between positive sexual selection and purifying natural selection affecting DNA sequences and gene expression is likely to differ between different stages of spermatogenesis (Figure 1). Genes expressed during the two first stages affect basic cell divisions and should be subject to strong purifying natural selection that prevents the spread of detrimental mutations^28,29^ and the appearance of disruptive selfish elements such as transposons^30^ and meiotic drivers^31^. The first stage, where there is a continuous mitotic proliferation of spermatogonia, is known to be very sensitive to mutations given the high number of mitotic cell divisions^32^. This aligns with the presence of strong purifying selection on both protein-coding genes and their regulatory elements that would benefit both spermatogenesis in males and cell proliferation and oogenesis in females (Figure 1). The genes underlying these early, basic stages in gamete production could potentially be subject to sex-specific selection since male fitness relies on fast production of many gametes (i.e., stronger positive sexual selection) while female fitness is ensured through fewer gametes of higher quality (i.e., stronger purifying selection). However, sex-specific gene regulation provides resolutions to such intra-locus sexual conflicts by allowing the two sexes to express different optimal phenotypes^33^. While the mitotic divisions of the female germ cells only occur in the embryonic gonads and then enter meiosis during fetal development and stay arrested, mitotic proliferation continues postnatal in males and gametes are produced throughout life.

**Figure 1.**
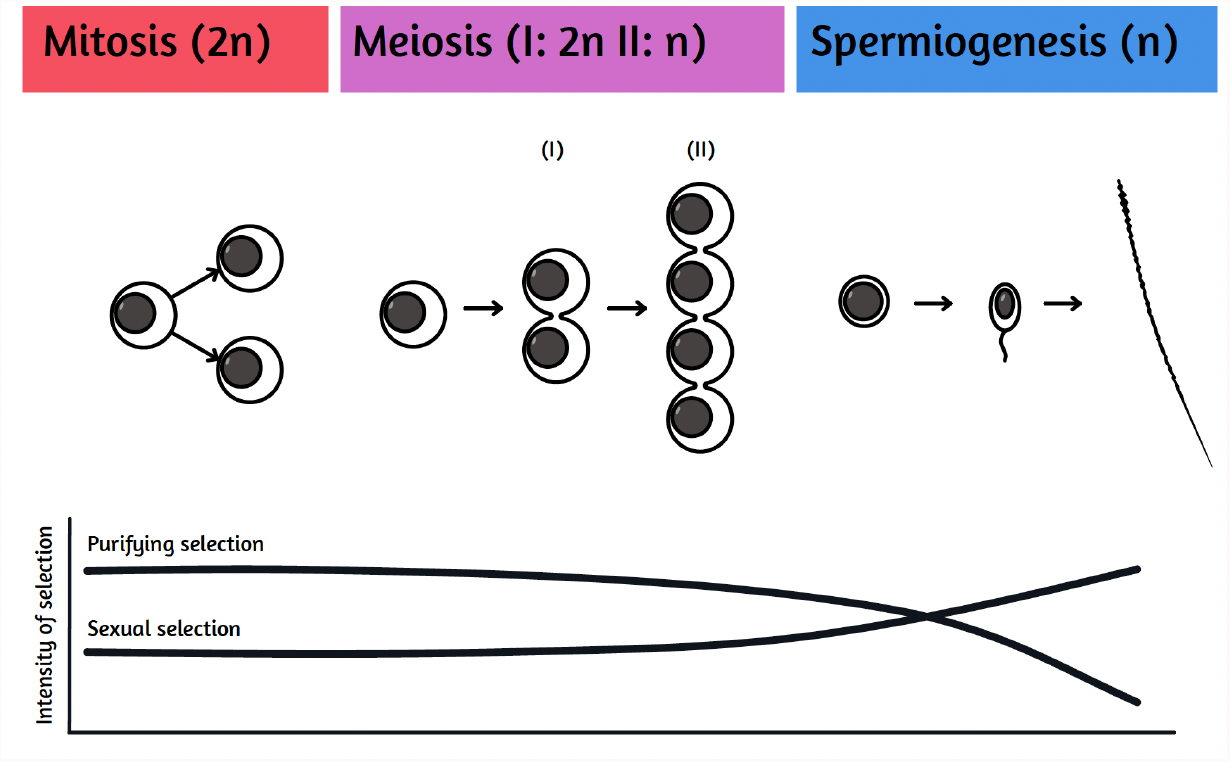
Expected balance between positive sexual selection and purifying natural selection across the different stages of spermatogenesis. Spermatogenesis can for simplicity be dived into three stages. The first stage includes several types of spermatogonia, where some cells are in constant mitotic activity to preserve the basal number of cells in the spermatogonia population while others start to mature and get ready for the next step. The second stage includes all cells that are in the different stages of meiosis also known as spermatocytes. The third stage includes the haploid spermatids (round spermatids and elongating spermatids). At this final stage, genes coding for sperm traits that will define characteristics such as swimming ability, velocity and compatibility with female reproductive fluid and female gametes, are expressed. Stages one and two are suggested to be subject of strict control ensuring the correct respective cell divisions that keep the whole process functional. By contrast, purifying selection on genes acting during the last stage of spermatogenesis is suggested to be relatively relaxed and sexual selection acting on sperm traits that affect success in sperm competition and/or are favored by cryptic female choice to be strong.

Spermiogenesis, the final stage of spermatogenesis should be more species-specific than the previous stages. We expect purifying selection on genes expressed during this final stage to be relatively relaxed and sexual selection to be strong in the form of both sperm competition and cryptic female choice^34^ (Fig 1). Because strong sexual selection acting on males may favor sperm traits in males that are detrimental for female fitness^35^, inter-locus sexual conflict could arise and speed up diversification^36,37^. With the increased use of single-cell RNA (scRNA) sequencing, we now have a better understanding of spermatogenesis, especially in mammalian and insect model organisms, all of them having XY sex determining systems^13,38–42^. This has allowed the identification of major germ and somatic cell types in the testis as well as the characterization of expression patterns throughout the different stages of spermatogenesis. It has thereby also become possible to study evolutionary aspects of spermatogenesis in greater detail. For example, comparisons between human, macaque and mouse showed conserved gene expression of spermatogonia and more diversification in the spermatid differentiation stage, which is consistent with relaxed purifying selection and/or increased sexual selection acting toward the later stages of spermatogenesis^39,40^. Taken together, previous theoretical and empirical studies in mammals and insects suggest faster adaptive evolution at the last stage of spermatogenesis.

The evolution of distinct male and female reproductive features are tightly intertwined with the evolution of sex chromosomes. A striking characteristics of mammalian spermatogenesis is that sex-chromosomes are inactivated during meiosis^43^ and their transcription is repressed at the post-meiotic stage^34^. Comparisons between human, macaque and mouse have revealed consistency in sex-chromosome silencing during meiosis (MSCI), except for some escape genes in the case of primates^39,40^. Apart from a few genes that remain repressed post-meiotically, many X-linked testis-specific genes are highly expressed during this very last stages of spermatogenesis (spermiogenesis). These genes evolve fast and tend to vary also among closely related species and may often be underlying the evolution of hybrid male sterility^34,44^.

Bird spermatogenesis, where spermatogenesis happens in the homogametic sex, is still poorly understood^45^, especially in passerines and has only been described morphologically^45,46^ or using bulk RNA sequencing^11^. However, in order to understand the different stages of spermatogenesis, cell diversity composition and underlying evolutionary processes, a single cell approach is necessary^10,38,42^. This is particularly important for amniotes where the testis is composed of seminiferous tubules and multiple cell types and cell stages are mixed^47^.

While the general expectations for the balance between evolutionary forces acting on genes expressed during spermatogenesis are similar for birds and mammals, there may also be some specific constraints derived from the different types of sex determination systems. The expectation of relaxed purifying selection and increased sexual selection acting toward the later stages of spermatogenesis^39,40^ is similar (Figure 1). However, since meiotic sex-chromosome inactivation (MSCI) is triggered by the absence of homologous sex-chromosome pairing partners in the heterogametic sex^48^, it does not apply to spermatogenesis in birds since males are the homogametic sex. Therefore, Z-linked testis-specific genes may be active during all stages of spermatogenesis in birds.

In this study we present a detailed molecular characterization of passerine spermatogenesis based on scRNA-seq of the testis of collared (*Ficedula albicollis*) and pied (*Ficedula hipoleuca*) flycatchers with the aim to reveal conserved and species-specific evolutionary features of this central process. The two closely related species of flycatchers are widely used in studies on ecology and evolution^49^ and male hybrids resulting from crosses between the two species are known to experience impaired sperm production^50^. There is also evidence for post-mating, pre-zygotic isolation^51^. These findings imply divergence in genes underlying spermatogenesis and make it possible for us to investigate conserved and diverged evolutionary features on a relatively short evolutionary time scale. We investigate the hypothesis of strongly conserved patterns of gene expression during the first stages of spermatogenesis and diverged features at the final stage in a ZW sex-determining system. Since our study is the first characterization of spermatogenesis at a single cell level for a ZW sex-determining system, an additional major goal is to assess the activity of the Z chromosome at the different stages of the process. We will also investigate whether Z-linked, testis-specific genes are more conserved or diverged at the different stages of spermatogenesis compared to testis-specific genes located on the autosomes.

## Results

### Characterization of bird spermatogenesis

Using the 10X genomics Chromium Single cell platform, we generated scRNA data from testis cell suspensions of two pied flycatchers and one collared flycatcher. After data pre-processing and cleaning, we recovered 4936 cells across our three samples that resulted in a consensus of 18 distinct clusters (Fig 2A) where each cluster contained cells from all three individuals (Fig 2B). We identified gene markers for each cluster, and a subset of the gene markers is shown in (Fig 2C, 2F) (Table S1). Most of the top cluster marker genes for flycatchers are not functionally annotated or do not have a corresponding ortholog in mammals. However, 65 representative flycatcher markers had unique orthologs in human and we found three genes that are also highlighted as important markers of different stages of spermatogenesis in mammals. A marker gene for undifferentiated spermatogonia, UCHL1, a pachytene marker, MYBL1, and an elongating spermatid marker, TSSK6 were found in mouse, human, macaque, and flycatchers. We also found several markers for mammalian spermatogenesis cell types that were expressed in our clusters of flycatcher spermatogenesis, (Fig S1, S2) and therefore were useful in the assessment of the identity of most of our clusters. Two of our 18 clusters were characterized as belonging to the initial stage of spermatogenesis (i.e. as spermatogonia), five to meiosis stages (i.e. spermatocytes) and six to the final stages of differentiating spermatids. In addition, five clusters were identified as somatic cells (Fig 2C). Among the markers indicative of clusters belonging to the somatic cells, we found mammalian markers for macrophages, sertoli cells, leydig cells and structural cells (Fig S2). Gene Ontology (GO) analysis based on the gene markers showed relevant biological processes for all the three main stages of spermatogenesis (Fig 2D). Consistent with this functional identification of clusters, velocity analysis and the PAGA graph obtained with ScVelo indicated a congruent order of the clusters with the main stages already identified (Fig 3). A general comparison of all markers from the three main stages with all the markers identified by Hermann et al., 2018 for mouse and humans, shows that the differentiating spermatids have significantly less shared markers between mammals and flycatchers than the first two stages (X^2^ -test with df = 2, X^2^ = 76.4, *p*-value < 2.2e^-16^) (Table S2, S3) (Fig 4).

**Figure 2.**
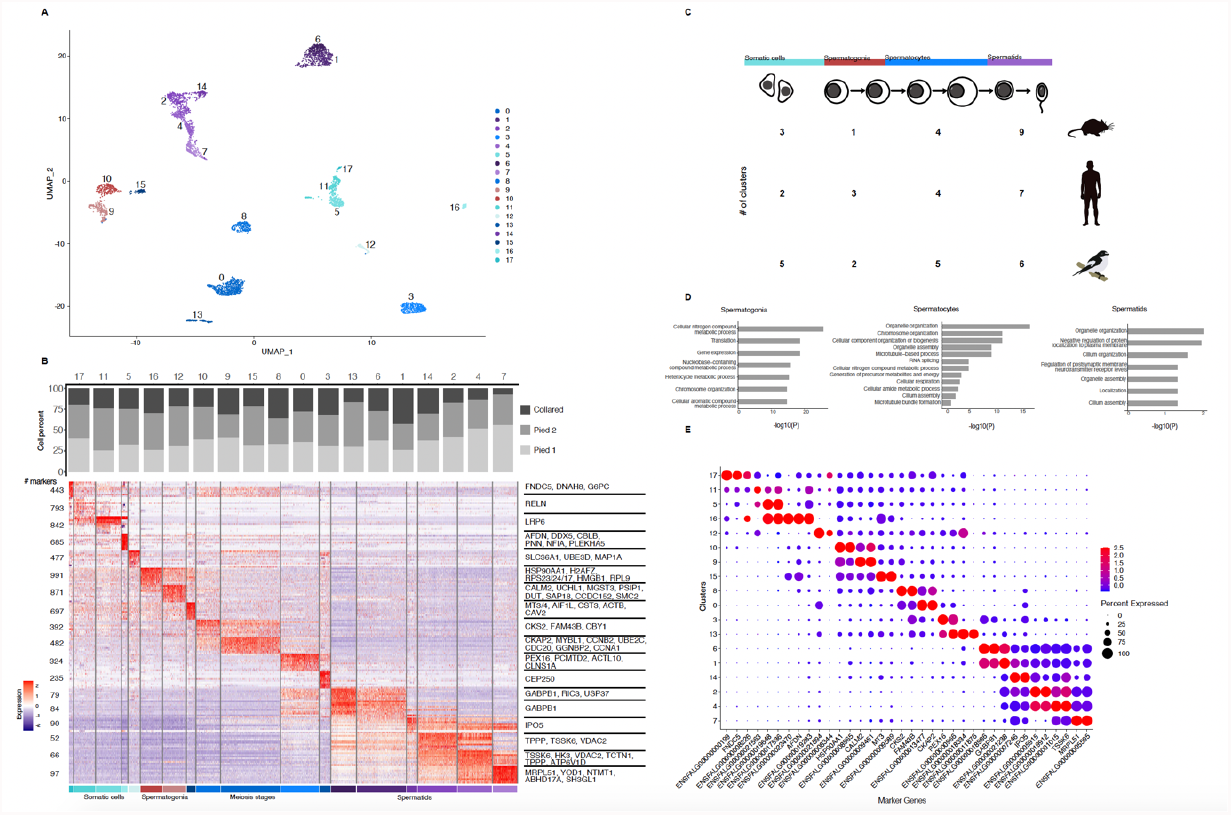
Single cell RNA sequencing of testis samples from Ficedula flycatchers reveals 18 spermatogenesis cell types. (A) Visualization of consensus clustering of testis cells from 3 flycatcher individuals in UMAP (Uniform Manifold Approximation and Projection). (B) Spermatogenesis cell cluster composition in terms of the percentage of cells from each testis sample (i.e. individual bird) and heatmap showing the mean expression of each top 10 marker genes of the 18 cell types found. Representative makers with annotated function are listed to the right and the total number of markers for each cluster are listed to the left side. (C) Comparison of the number of clusters of testis cell types found in flycatchers, human and mouse using a similar methodology (data from Hermann et al. 2018). (D) Representative GO terms for the three main stages of spermatogenesis in flycatchers, with FDR-corrected p-values shown for each GO term. (E) Subset of the best 2 or 3 markers specific to flycatchers for each cluster, most of the markers lack functional annotation or do not have an ortholog and therefore are shown with their Ensembl gene ID.

**Figure 3.**
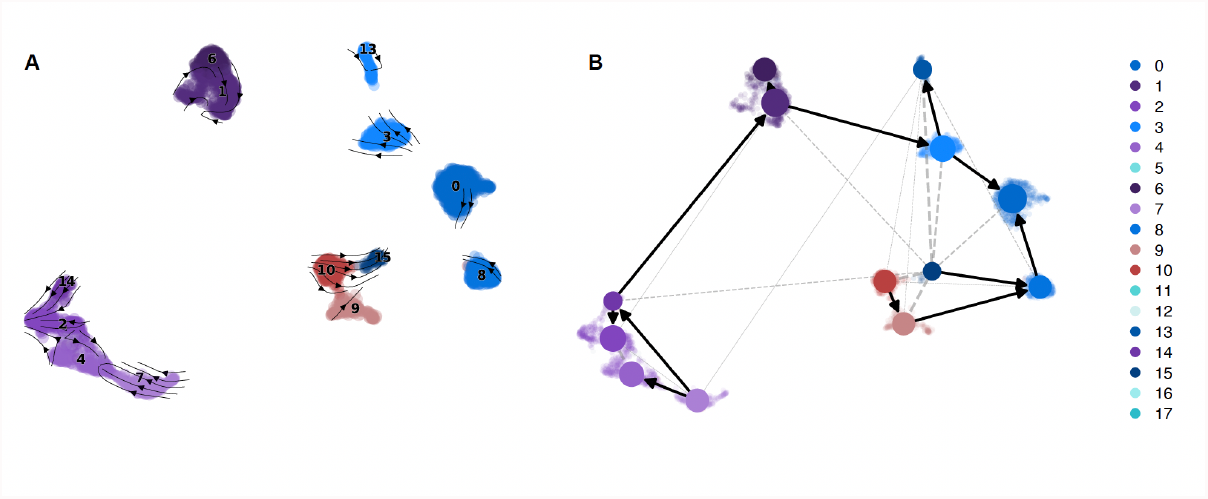
Trajectory of Ficedula flycatcher testis cells. (A) Velocity dynamics of germline cell clusters. (B) PAGA graph of germline cell clusters, the arrows indicate connection strength and possible direction of differentiation.

**Figure 4.**
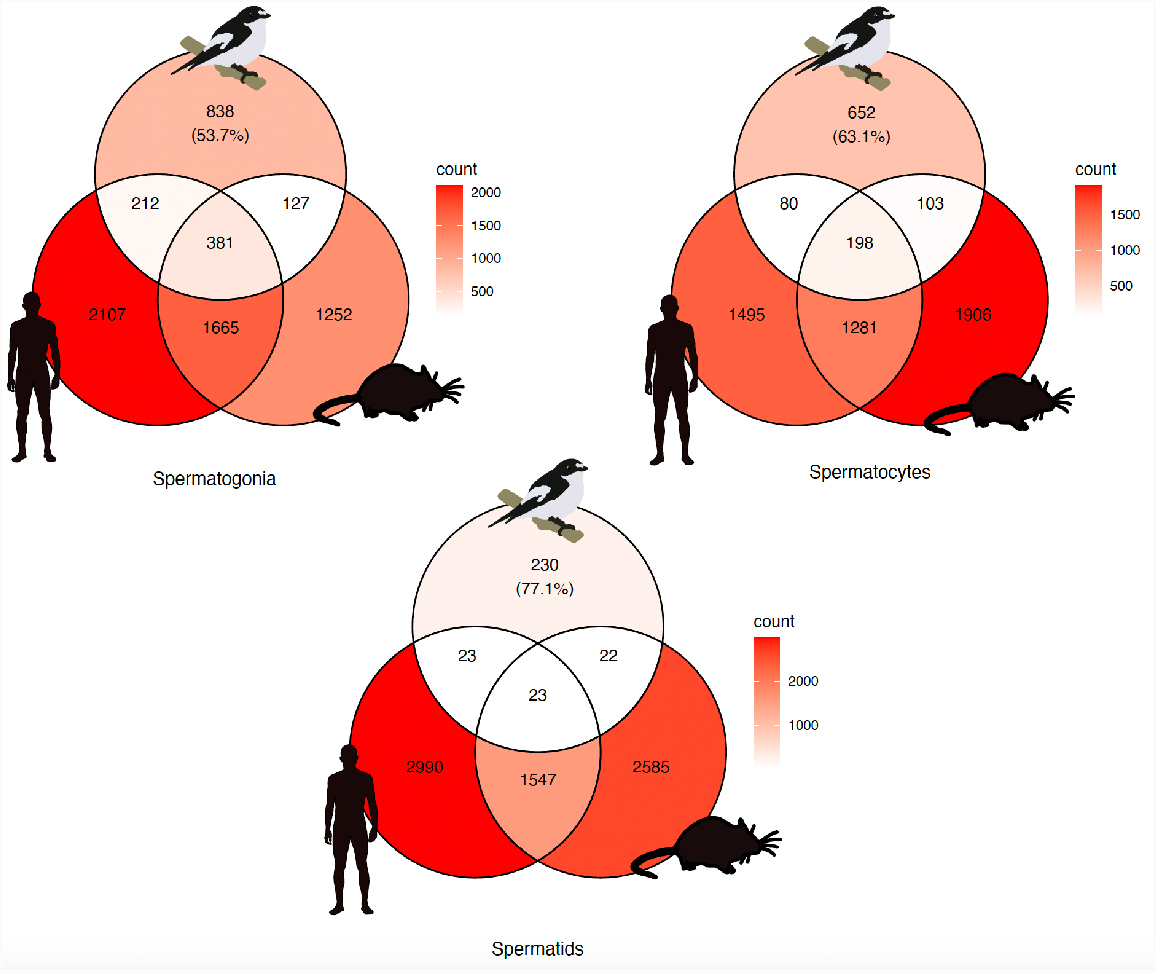
Venn diagrams for the three main stages of spermatogenesis showing shared and private marker genes between flycatchers, human and mouse. The percentage in brackets corresponds to the percentage of private flycatcher markers. Spermatids share the least number of ortholog markers between flycatchers and either human or mouse as compared to the other stages of spermatogenesis.

### Gene expression patterns

Excluding the somatic cell clusters and considering all expressed autosomal and Z-linked genes, we found a similar mean expression across the cell clusters (Fig 5A) (Table S4). At this last stage of spermatogenesis when spermatids are differentiating, the increase in variance in gene expression is due to a very high expression of a few genes. Cluster 13 belonging to the meiosis stages, is the only cluster showing a pronounced tendency for a higher average expression of Z-linked genes with respect to the autosomal genes. When restricting the analysis to the top 500 expressed genes, we still find the pattern described above for cluster 13, and that there is an increase in mean expression along the ordered timeline (Fig 5B).

**Figure 5.**
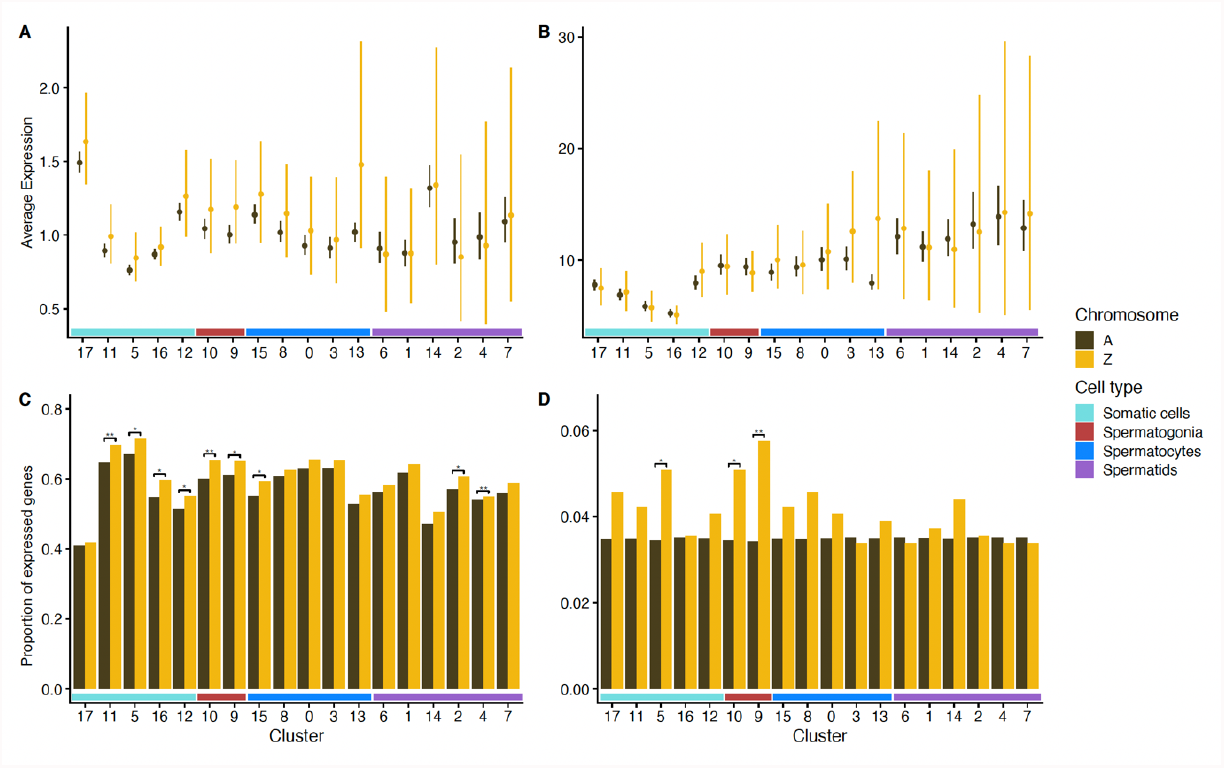
Average expression and proportion of expressed genes across all 3 testis samples from flycatchers for each cell clusters. Average expression of all genes located on the autosomes or on the Z chromosome (A) For all expressed genes and (B) for the top 500 expressed autosomal and Z-linked genes. Error bars were obtained by bootstrapping. (C) There is a significant enrichment of Z-linked genes expressed in four somatic clusters, two spermatogonia clusters, one meiosis cluster and two spermatid clusters. For all genes with expression level >0.01. (D) There is a significant enrichment of Z-linked genes among the top 500 genes expressed in one somatic cluster, and in two spermatogonia clusters.

We also tested for an enrichment of Z-linked genes among the expressed genes in each cluster. When considering all genes with average expression > 0.01, we found a general trend of enrichment of Z-linked genes across all stages of spermatogenesis, but significant enrichment of Z-linked genes was only found for genes expressed in four somatic clusters, the two spermatogonia clusters, one meiosis cluster and two spermatid clusters (Fig 5C) (Table S5). This finding remains consistent for the two spermatogonia clusters and one somatic cluster when restricting the analysis to the top 500 genes (Fig 5D) (Table S6).

We calculated fold-change of gene expression for each cell cluster between the two flycatcher species using Wilcoxon rank sum test for differential expression (DE) analysis (Fig S3-S6) (Table S7). Very few or no genes were differentially expressed in the 5 clusters of somatic cells and in the two clusters of spermatogonia cells. Most DE genes were found in two clusters belonging to the meiosis stage and in five clusters belonging to the last stage of spermatogenesis. Of these, cluster 1 had the most DE genes of all clusters (i.e. 291 DE genes) (Fig 6A). Consistent with findings in species with XY sex determination, we thus find most differences in gene expression between the two species towards the later stages of spermatogenesis (generalized linear model with binomial distribution, 1.63 ± 0.07, Z = 25.14, *p*-value < 2e^-16^) (Table S8). We tested for overrepresentation of Z-linked DE genes and found that one of the cell clusters going through meiosis (i.e. cluster zero) had a significant overrepresentation of DE genes on the Z chromosome (hypergeometric test, p=0.006) (Table 1). We lack statistical power to test for overrepresentation of Z-linked genes in cell clusters with few DE genes but most clusters with several DE genes show a tendency for overrepresentation of Z-linked genes (Fig 6A). We found significant GO terms among the DE genes in clusters belonging to meiosis and the final stage of spermatogenesis (i.e. cluster 0, 3, 6, 1, 2 and 4) (Table S9). These GO terms were mainly related to cellular respiration and cell motility. Moreover, we found eleven DE genes located on the mitochondrial chromosome, four autosomal genes that coded for mitochondrial proteins and two genes on the Z chromosome that also coded for mitochondrial genes (Table 2).

**Table 1.** Numbers of differentially expressed (DE) genes and non-differentially expressed (non-DE) genes between collared and pied flycatchers per cluster for autosome genes and Z-linked genes. Mitochondrial DE genes and DE genes with unknown position in the genome were excluded from this particular analysis.

| Stage | Cluster | Autosomes |  | Z Chromosome |  | P-value |
| --- | --- | --- | --- | --- | --- | --- |
|  |  | DE | non DE | DE | non DE |  |
| Somatic cells | 17 | 0 | 2508 | 0 | 113 | 1 |
| Somatic cells | 11 | 0 | 689 | 0 | 43 | 1 |
| Somatic cells | 5 | 0 | 693 | 0 | 37 | 1 |
| Somatic cells | 16 | 0 | 2352 | 0 | 133 | 1 |
| Somatic cells | 12 | 0 | 1235 | 0 | 68 | 1 |
| Spermatogonia | 10 | 1 | 929 | 0 | 43 | 1 |
| Spermatogonia | 9 | 2 | 818 | 0 | 40 | 1 |
| Meiosis stages | 15 | 0 | 1852 | 0 | 83 | 1 |
| Meiosis stages | 13 | 0 | 1759 | 0 | 85 | 1 |
| Meiosis stages | 0 | 36 | 354 | 8 | 12 | **0.00044** |
| Meiosis stages | 8 | 5 | 767 | 0 | 35 | 1 |
| Meiosis stages | 3 | 39 | 541 | 4 | 35 | 0.2827 |
| Spermatids | 6 | 104 | 348 | 9 | 16 | 0.1092 |
| Spermatids | 1 | 215 | 397 | 12 | 23 | 0.6057 |
| Spermatids | 14 | 1 | 993 | 0 | 46 | 1 |
| Spermatids | 2 | 44 | 657 | 6 | 41 | 0.085 |
| Spermatids | 4 | 32 | 497 | 4 | 26 | 0.119 |
| Spermatids | 7 | 19 | 1160 | 1 | 50 | 0.5742 |

**Table 2.**
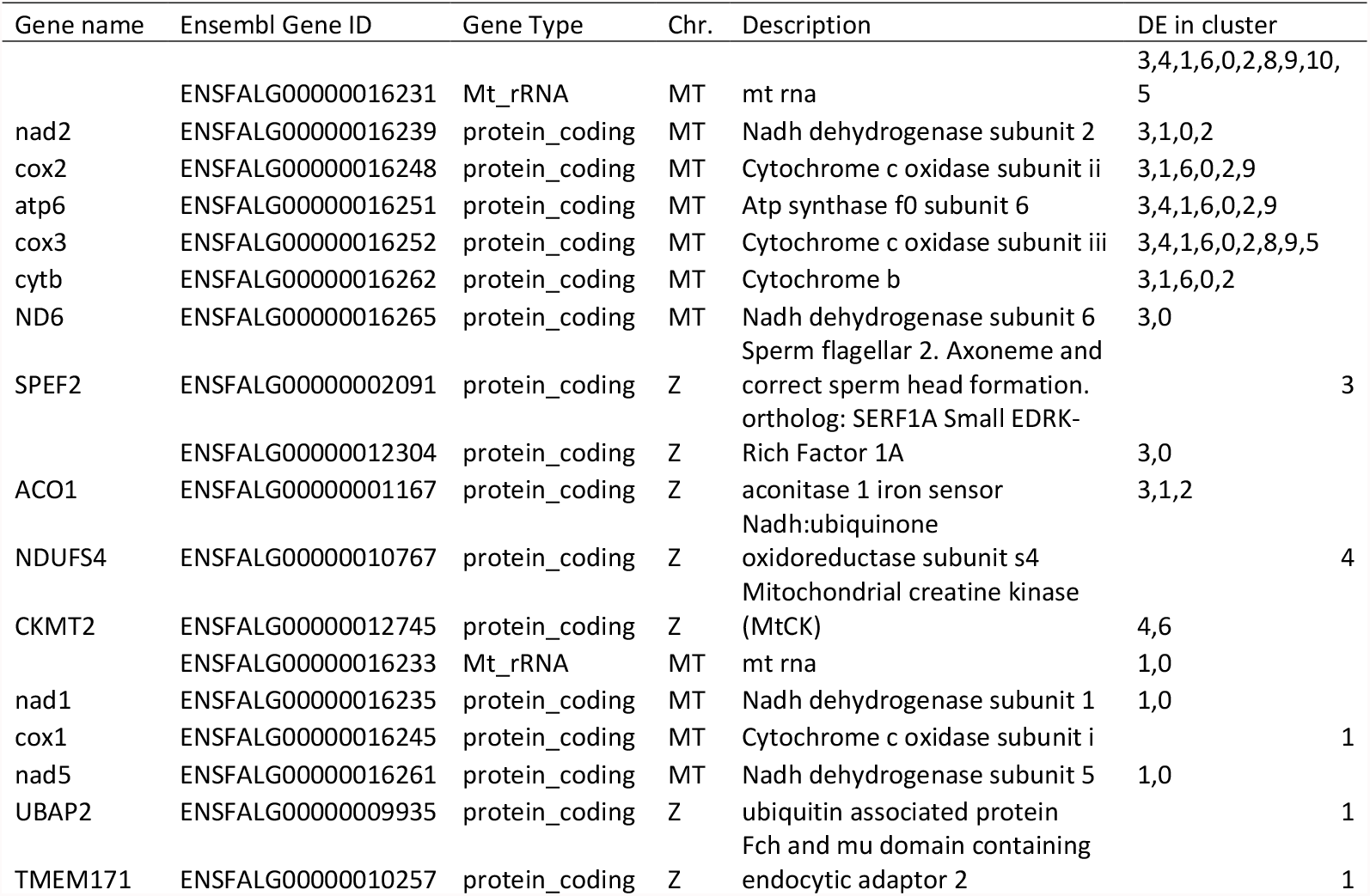
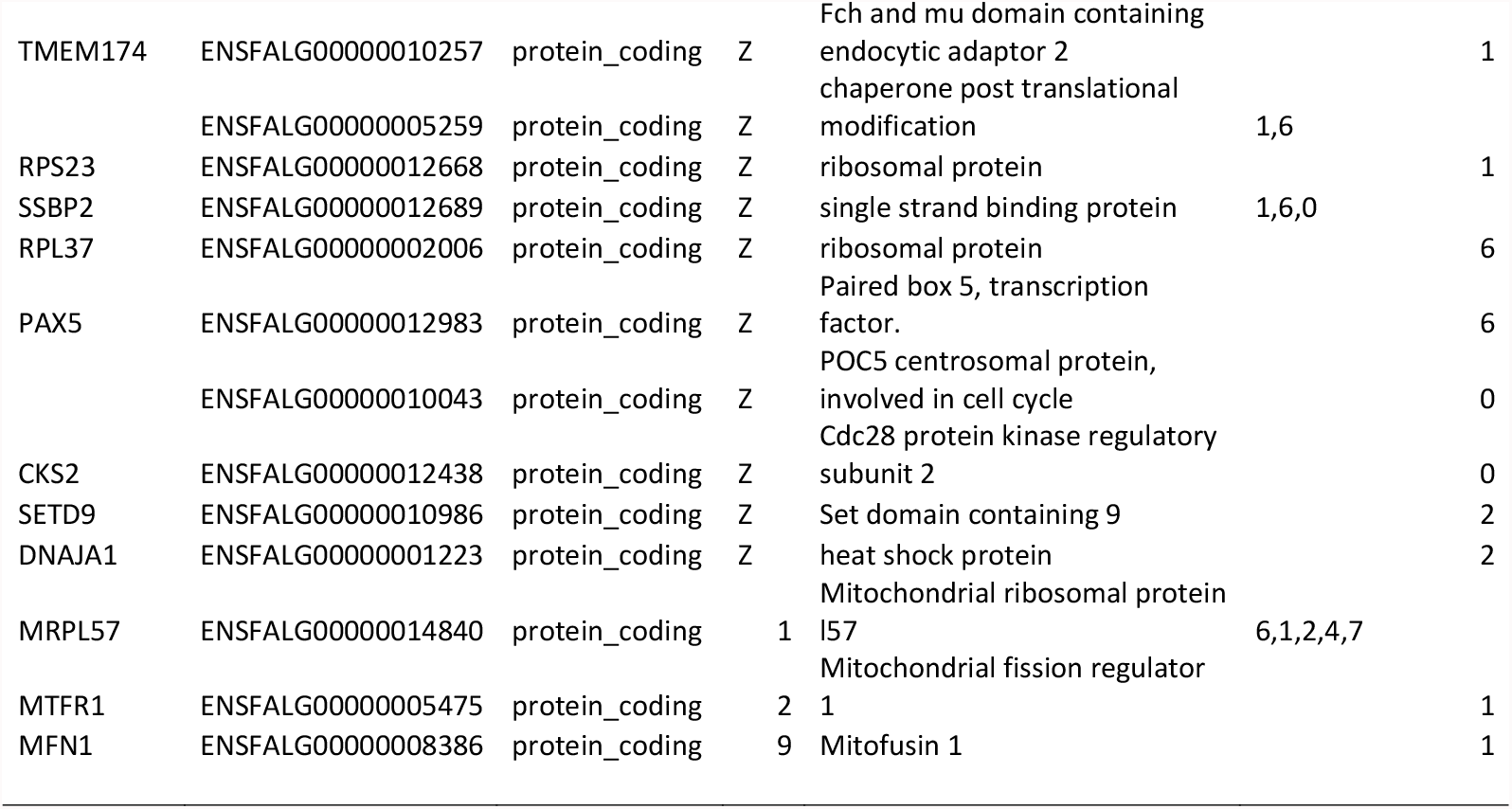
DE genes related to cellular respiration and cell motility. Subset of DE genes between collared and pied flycatchers located on the mitochondrial chromosome and on either the Z-chromosome or autosomes but coding for mitochondrial proteins.

**Figure 6.**
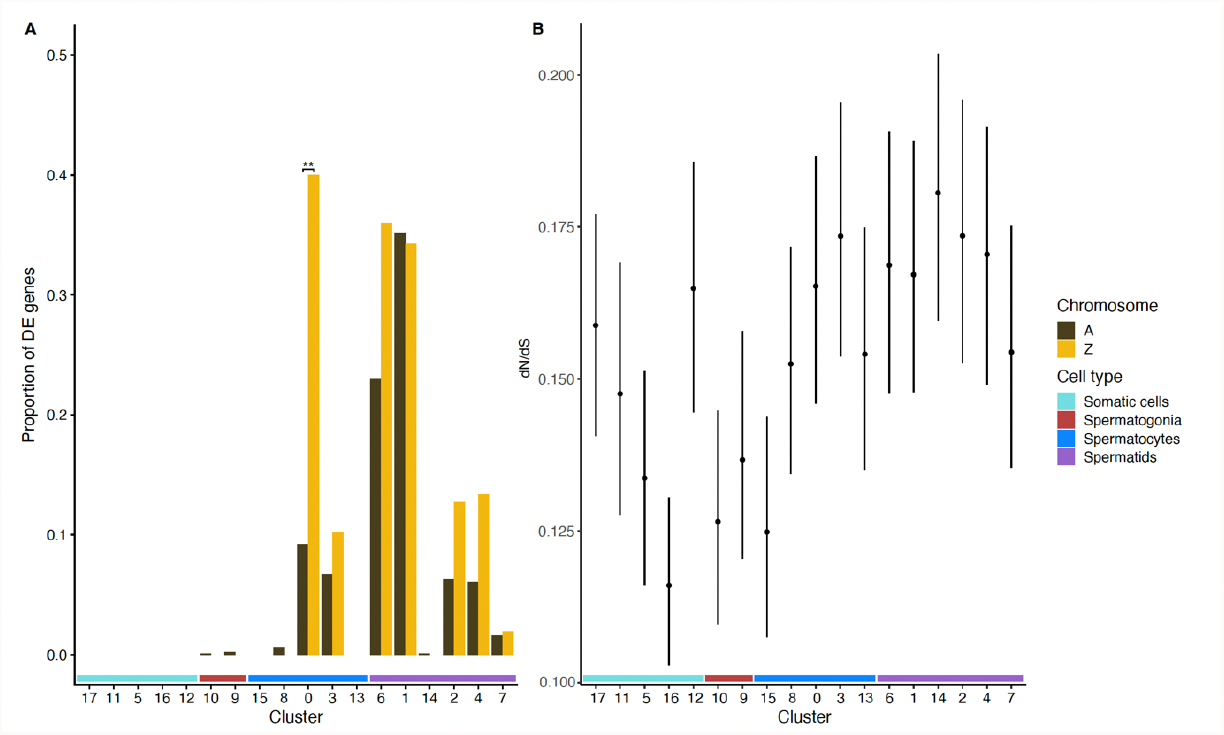
Patterns of differential gene expression between the two flycatcher species among the different testis cell clusters and signatures of gene sequence evolution. (A) Proportion of DE genes between collared and pied flycatchers in each cluster. DE genes are only found in clusters belonging to spermatogenesis, most of them are present in the meiosis clusters and the spermiogenesis clusters. There is a general tendency for overrepresentation of Z-linked genes among the DE genes between the two species. However, only spermatocytes of Cluster 0 belonging to the meiosis stage show a significant overrepresentation of Z-linked genes. (B) d_N_/d_S_ mean value for the top 500 expressed genes by cluster. There is an increase in d_N_/d_S_ mean value along the spermatogenesis timeline consistent with a higher constraint at the beginning of the process and relaxed constraint towards the end.

### Signatures of gene sequence evolution across the stages of spermatogenesis

The average *d*_N_/*d*_S_ values between collared and pied flycatchers for the genes expressed in the different cell clusters ranged from 0.1 to 0.2, which indicates that genes are on average evolving under constraint across all stages of spermatogenesis. However, the values are generally smaller, which is indicative of more constraint, for the somatic and for the first cell clusters, while the *d*_N_/*d*_S_ values increase towards the end of the timeline, which is indicative of less constraint or an increasing contribution of positive selection (Figure 6B). These findings are hence indicative of stronger constraint on the initial spermatogonia stages of spermatogenesis and that the magnitude of purifying selection slightly relaxes toward the end of the timeline, or that that there is positive selection acting on an increasing fraction of sites within the genes that are expressed during the final stages of spermatogenesis.

## Discussion

Functional spermatogenesis is a pre-requisite for male fertility in sexually reproducing organisms, but has previously only been well characterized in a handful of model systems such as *Drosophila* and a few mammalian species^13,38–40,42^. In all these previously studied model systems males constitute the heterogametic sex, implying that some major evolutionary features are shared between them such as the regulatory constraints associated with meiotic sex chromosome inactivation (MSCI) and post meiotic gene repression, which select against sex-linked meiotic genes. Based on scRNA-seq of the testis of two closely related passerine bird species, where males constitute the homogametic sex, we can reveal both, striking evolutionary similarities and differences, compared to previous findings based on the study systems with heterogametic males. One of our major goals was to characterize the relative average expression of Z-linked genes to autosomal genes along the different stages of spermatogenesis. We know from oogenesis in XY systems, which happens in the homogametic sex, that there is an absence of MSCI given that there are no unpaired chromosomes. However, before the moment of chromosome synapsis one of the normally silenced X chromosomes needs to be unsilenced causing a hyperexpression of the X chromosome (i.e. dosage decompensation)^52^. Birds have been found to have partial dosage compensation regulated on a gene-by-gene basis^53,54^. This is mainly evident in the heterogametic females where the Z:A ratio has been found to be < 1 while homogametic males have a Z-linked average expression similar to the autosomes (i.e. Z:A ∼ 1).

Nevertheless, we observe a high average expression of the Z chromosome with respect to the autosomes in one of the cell clusters belonging to meiosis (cluster 13; Fig 5A, 5B). This signal appears very similar to the moment in mammal oogenesis when the X chromosome is showing dosage decompensation during meiosis to enable the following chromosome synapsis^52^. Since we know that some genes on the Z chromosome show dosage compensation while others do not, it is possible that the signal we observe reflects decompensation of the compensated genes similarly to what happens during mammalian oogenesis.

In agreement with earlier studies investigating signals of molecular evolution across different stages of spermatogenesis in mammals^34,39,40,55^, we find that the first stage of spermatogenesis shows several signs of strong evolutionary constraint. These signs include higher conservation of gene expression between the two flycatcher species at this stage compared to the later stages (Fig 6A), higher conservation of protein-coding sequences of genes expressed at this stage (Fig 6B), and more detected shared orthologous gene markers between flycatchers and mammals among the gene markers used to characterize the cell clusters (Fig 4). Like in mammals, we also find a significant enrichment of Z-linked genes in the two clusters of this stage^55^. All together these results reaffirm the prediction that basic processes such as mitotic cell division are highly conserved even among divergent taxa due to strong purifying selection to keep functionality.

During the second stage of spermatogenesis when the different steps of meiosis occur, we expected to find lesser constraint than the first stage of spermatogenesis. However, we find evidence of even lesser constraint than expected based on earlier observations in XY systems. Consistent with expectations, genes expressed in the second stage show lower conservation of protein-coding sequences compared to genes expressed in the first stage of spermatogenesis. However, two cell clusters belonging to the stages of meiosis contain an unexpected high proportion of genes that are differentially expressed between the two flycatcher species (Fig 6A). This result indicates that gene expression at this stage is less constraint by purifying natural selection and/or by intra-locus sexual conflict as compared to mammals. While this finding may be specific to the flycatcher system, given the absence of MSCI during male meiosis in ZW systems it could also be shared by other ZW species. In mammals, MSCI and post-meiotic repression of X-linked genes result in many testis-biased genes having been copied from the X chromosome to the autosomes.^56–58^. Alternatively, X-linked testis-biased genes are highly expressed in earlier stages of spermatogenesis and the transcripts prevail until the final stage where its function is necessary^59^. By contrast, testis-biased genes can remain on the Z chromosome and be synthetized and expressed when needed in birds. We indeed find a significant signal of enrichment of Z-linked genes among the genes that were differentially expressed between the two flycatcher species during this second stage of spermatogenesis. While this result again may be specific for the flycatcher case, it is also consistent with relaxed constraint specifically associated with the absence of MSCI which is a universal feature for birds and all ZW systems. Studies of additional species, where spermatogenesis happens in the homogametic sex, are needed to provide a broader picture. Nevertheless, since male fitness relies on fast production of many gametes, we suggest that this relief of constraint may potentially explain why bird spermatogenesis is four times faster compared to spermatogenesis in mammals.

As predicted, the third and last stage of spermatogenesis in the flycatchers shows even more clear signs consistent with relaxed purifying natural selection and/or strong positive sexual selection than the second stage. Protein-coding sequence divergence of genes expressed in the last stage is on average highest among all stages and we find a significantly higher proportion of DE genes between the two flycatcher species both in terms of number of DE genes and in terms of number of cell clusters with DE genes. In agreement with what has been described for mammals we also find more private markers at this stage and overall, less genes, for which one-to-one orthologs to mammals could be identified. One possible explanation for this observation is the presence of more de novo genes that are species-specific at this stage, similar to what has been found among macaque, human and mouse^40^. During this last stage of spermatogenesis, when the sperm cells obtain traits that influence swimming ability and other important functions known to affect success in sperm competition and cryptic female choice, a relatively high degree of expression of sex chromosome linked genes is expected^34^. This is because sex chromosomes are in general known to accumulate genes with sex-biased functions^60^. Since the Z-chromosome spends most of its time in males, the fixation of Z-linked mutations with male-biased fitness functions are expected to be favored by positive sexual selection^61–64^. There were tendencies for enrichment of expressed Z-linked genes both among all the clusters at this stage and among the DE genes between the two species of flycatchers. These findings are consistent with evidence of an overall elevated sequence and expression divergence of Z-linked genes compared to the autosomal genes in flycatchers^11,65^ and in other birds^63,66^.

However, fast evolution of Z-linked genes can aside from positive sexual selection also be caused by genetic drift in combination with a higher mutation rate in males^67,68^. The observed tendency for Z-chromosome enrichment of expression divergence associated with the last stage of spermatogenesis could therefore also reflect relaxed purifying selection instead of increased positive sexual selection. However, across the two last stages of spermatogenesis, we find several significant GO terms for biological processes among the DE genes between the two flycatcher species that are mainly related to cellular respiration and cell motility. The presence of GO terms related to cell motility reinforces the idea that sexual selection related to sperm performance has an important effect on the fast evolution of genes coding for these traits at the final stage of spermatogenesis. Since F1 hybrids resulting from crosses between collared and pied flycatchers experience hybrid dysfunction in terms of severely reduced fertility^50^ and in terms of elevated metabolic rate^69^, our results add to previous evidence suggesting that mito-nuclear incompatibilities may be causing or contributing to such dysfunction.

There are some major similarities between XY and ZW systems in terms of the selection pressures acting at the different stages of spermatogenesis with strong purifying selection during the first stage of spermatogenesis and sexual selection or relaxed purifying selection during the final stage. However, a major difference is the developmental constraint that heterogametic males experience in XY systems that males in ZW systems lack. This lack of constraint in theory allows the Z chromosome to keep testis-biased genes that also may include genes with sexually antagonistic fitness effects. We detect signals of fast evolving Z-linked genes expressed not only during the last stage of spermatogenesis but also already at the second stage thereby revealing a possible key evolutionary difference between this process in birds and mammals. We suggest that the high throughput of bird spermatogenesis, which is four times faster compared to mammals^70^, at least partly is explained by relief of evolutionary constraint, possibly also connected to an advantage in the intra-locus sexual conflict considering the optimal solution to the trade-off between gamete number and quality. Our study is the first characterization of spermatogenesis at a single cell level for a ZW system and it thereby opens the doors for future studies exploring various aspects of spermatogenesis in ZW systems such as infertility, molecular evolution and sex-chromosome evolution.

## Methods

### Sample collection, cell suspension preparation and sequencing

Two pied flycatcher (*Ficedula hypoleuca*) and one collared flycatcher (*Ficedula albicollis*) males were sampled from the monitored populations on Öland (57°100N, 16°580E), Sweden^49^ in May of 2019. The individuals were trapped in nest boxes while defending territories before nest building at the beginning of the breeding season. These birds were briefly kept in outdoor aviaries and then transported in individual cages over night to our lab facilities at Uppsala University. By choosing this timing during the breeding season, we ensured that the individuals possessed fully functional testis, because the testis degenerates once the reproductive season is over in this species. The animals were sacrificed and immediately dissected. All animal handling was done following Swedish regulations with the required permits approved.

The testes were placed in a cold petri dish with PBD BSA buffer, cleaned from any other visible tissue cells and cut in half. A half of testis tissue was then put in 3ml of buffer PBD BSA and dissociated by mechanical dissociation using the gentleMACS Dissociator (Miltenyi Biotech, Bergisch Gladbach, Germany) with the gentleMACS C Tubes using the preset for mouse spleen as recommended by the manufacturer. Next, the cell suspension was centrifuged to assure all cells would get down the cap and the walls of the tube. Finally, the cell suspension was homogenized by pipetting and filtered using a cell strainer of 70(μm).

After mechanical dissociation, an aliquote of the cell suspension was stained with propidium iodide and Hoechst for live death cell staining, examined under the microscope, and cell viability was estimated to be at least 80%. Cells were counted using a Neubauer chamber and the cell suspension was then diluted to achieve the required concentration of 1 × 10^6 cell/ml. The final cell suspension of a total volume of 500 ul was immediately delivered to the sequencing platform for library preparation with 10X genomics Chromium Single Cell 3’ v3 kit for scRNAseq. The whole process between sacrificing the animals and handing in the final cell suspension, lasted 2 to 3 hours during which the samples were kept on ice at all times. The 3 libraries were sequenced in one NovaSeq SP flow cell, yielding an approximate of 215 million reads per sample.

### Data processing and analysis

We created a custom reference for cell counting using the 10x Genomics Cell Ranger v. 4.0.0.^71^ command mkref following default settings and the well documented CellRanger pipeline. For that, we used the publicly available genome .fasta file (v. 1.4) of the collared flycatcher and the collared flycatcher annotation .gtf file from *Ensembl* (v. 1.4). Count matrixes for every gene in each individual cell for the three samples were obtained. Using this output and the .gtf file, a .loom file was generated for each of the three samples using the command run10x from the python package Velocyto v. 0.17.17^72^. Finally, this file was exported to Seurat v. 3.2.0^73^ for filtering, normalization and clustering (described below).

### Filtering, normalization and clustering

The three .loom files were imported to R, transformed to Seurat objects and then merged to a single object. We filtered out all cells with less than 200 features and more than 2500 features as well as cells with less than 5% of mitochondrial genes. The normalization was done using anchors to integrate the 3 samples and SCTransform as documented in the Seurat pipeline. Finally, we followed the standard Seurat workflow for clustering using UMAP (Uniform Manifold Approximation and Projection). Because spermatogenesis is a continuous non-discrete process, we obtained a big cluster containing a heterogenous cell composition with no clear gene markers, therefore that cluster was excluded. The remaining cells were re-clustered.

Marker genes for each cluster were identified using the Seurat function FindAllMarkers. The default setting uses the Wilcoxon rank sum test and calculates average log fold change for each gene between the clusters. We also found all markers using the ROC standard AUC classifier method, most of which were the same as the markers found with Wilcoxon rank sum test. We selected the top 3 best markers for each cluster having the highest AUC value Figure 2 F.

### Characterization of the cell clusters

We searched for the presence of all the previously identified markers for the different spermatogenesis stages in human^38,41^, mouse^38,42^ and macaque^39,40^ in our flycatcher cell clusters. By crossing the information of the previously identified markers found in our data we assigned identities to the cell clusters to the four major groups of cells found in spermatogenesis: Spermatogonia cells, meiotic cells, differentiating spermatid cells and somatic cells.

### Gene Ontology analysis

We performed a Gene Ontology (GO) enrichment analysis for biological processes using the webtool ShinyGO v.0741^74^ with the collared flycatcher as a background. To remove redundant and/or nested GO terms we used the web tool REVIGO^75^. This analysis was done per cluster using all identified markers and afterwards per cluster using all the differentially expressed (DE) genes (see below).

### Velocity analysis

To confirm the identity of the clusters with another analysis and infer the most likely time trajectory, we used the python package scVelo^76^ to run velocity analysis, pseudotime analysis and a PAGA (Partition-based graph abstraction) graph^77^. With that purpose we exported our Seurat object to .h5dr format compatible with scVelo and followed their well-documented pipeline.

Once all our clusters were assigned to one of the main stages of spermatogenesis, we performed a comparison with mouse and human using one to one orthologs. We downloaded the complete list of markers found by Hermann et al. for mouse and human and we grouped the clusters in the three main stages of spermatogenesis and identified the shared markers among flycatchers, mouse and human. The shared markers were displayed using Venn plots.

### Gene expression patterns at different stages of spermatogenesis

We calculated average expression of all genes per cell cluster for our merged object using the AverageExpression function after using the LogNormalize function in the RNA counts matrix. Then we added as metadata the location of each gene, either on the autosomes or on the Z chromosomes. We subset the data to all genes expressed and to the top 500 expressed genes, respectively, and performed a bootstrap resampling with replacement to generate confidence intervals. To test for enrichment of Z-linkes genes in each cluster we performed a hypergeometric test for over-representation using the R function phyper and lower tail false.

### Molecular signatures of evolution throughout the cell clusters

The *d*_N_/*d*_S_ ratio per cluster was computed as the ratio of average *d*_N_ and average *d*_S_ for the top 500 expressed genes based on collared flycatcher specific *d*_N_ and *d*_S_ values^78^. We subset outlier genes having a *d*_S_ >1 and *d*_N_/*d*_S_ >2. Finally, we performed a bootstrap resampling to generate confidence intervals.

We used the metadata of species in our Seurat object to perform the comparison of gene expression at the different stages of spermatogenesis between the two species of flycatchers. The function FindMarkers was used to perform a Wilcoxon rank sum test to calculate average fold change and find DE genes between the two flycatcher species within each cluster. We used the adjusted *p*-value with a threshold of 0.05 for significance of average fold change. In order to visualize the DE genes, volcano plots were computed using the package enhanced volcano. We performed a hypergeometric test on the DE genes of each cluster to test for over-representation of Z-linked genes using the R function phyper. To test whether there was a significant difference of DE genes between the stages, we implemented a generalized linear model with binomial distribution of the response variable. We used the cbind function to consider DE genes or non-DE genes as binary response variable and we used spermatogenesis stage as explanatory variable. Finally, we performed a GO analysis on the DE genes using the same methods described above.

## Author Contributions

AQ and JCS conceived the study. AQ, JCS and CC collected the samples. JCS sacrificed the birds. CC performed the dissections. JCS, CC and CB adapted and carried out the cell suspension protocol. JCS and MS performed the single cell clustering and bioinformatics analysis. JCS, MS and CFM performed and discussed the molecular evolution and statistical analysis. JCS, MS, CFM and AQ discussed and interpreted all the results. JCS and AQ wrote the manuscript. All authors commented and approved the final version of the manuscript.

## Acknowledgements

We are thankful for help during the collection of the data and/or the development of the manuscript to William Jones, Katerina Guschanski, Tom van der Valk, David Weatcroft, Mahvash Jami, Javier Florenza García and all Qvarnström lab field assistants. We also thank the Microbial Single Cell Genomics Facility at SciLifeLab for help developing the cell suspension protocol and for letting us use their lab facilities. Sequencing was performed by the SNP&SEQ Technology Platform in Uppsala. The facility is part of the National Genomics Infrastructure (NGI) Sweden and Science for life Laboratory. The SNP&SEQ Platform is also supported by the Swedish Research Council and the Knut and Alice Wallenberg Foundation. Computations were performed on resources provided by the Swedish National Infrastructure for Computing (SNIC) at Uppsala Multidisciplinary Center for Advanced Computational Science (UPPMAX). Funding: This work was supported by the Swedish Research Council, grant number 2016–05138 and 2012–03722 to AQ. CFM is funded by grants to Hans Ellegren from the Swedish Research Council (2013/08271) and Knut and Alice Wallenberg Foundation. MS is funded by IUF (Institut Universitaire de France) and ANR 21 T-ERC PLEIOTROPY and received a stipend from Wenner-Gren for a sabbatical at Uppsala University.

## Ethical permits

Permit for keeping flycatchers in aviaries and sacrificing maximum 17 flycatchers per year. Swedish environmental protection agency Natur vårds verket (NV-01203-18) valid from 2018-05-01 till 2019-06-30.

## Data availability

All data and code will be available in a public repository (pending).

