## Supplementary material for "Single-Cell Transcriptomics reveals relaxed evolutionary constraint of spermatogenesis in two passerine birds as compared to mammals"

The supplementary material includes:

Fig S1-S6 and tables 2,3, 5,6,8. The rest will be available as separate files (pendant).

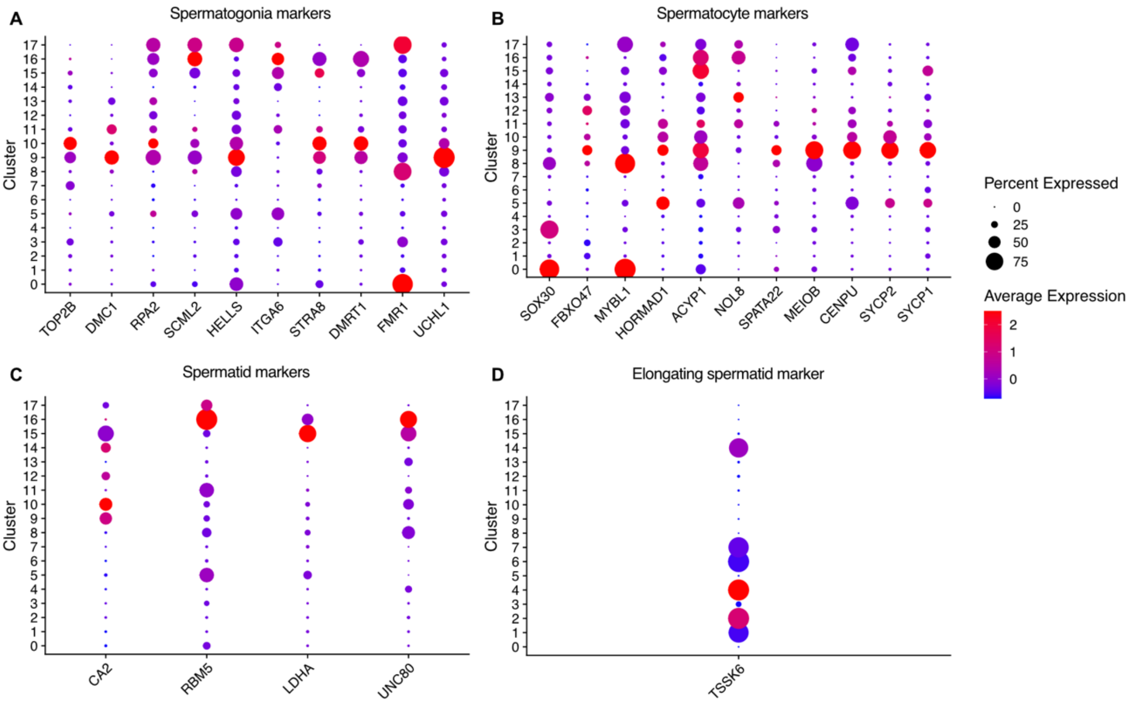


**Supplementary Figure 1. Average expression of mammalian testis marker genes in flycatcher tests cell clusters and the percentage of cells per cluster where each gene is expressed.** The Y-axis shows known mammalian gene markers for (A) spermatogonia, (B) spermatocytes, (C) spermatid, and (D) elongating spermatids.

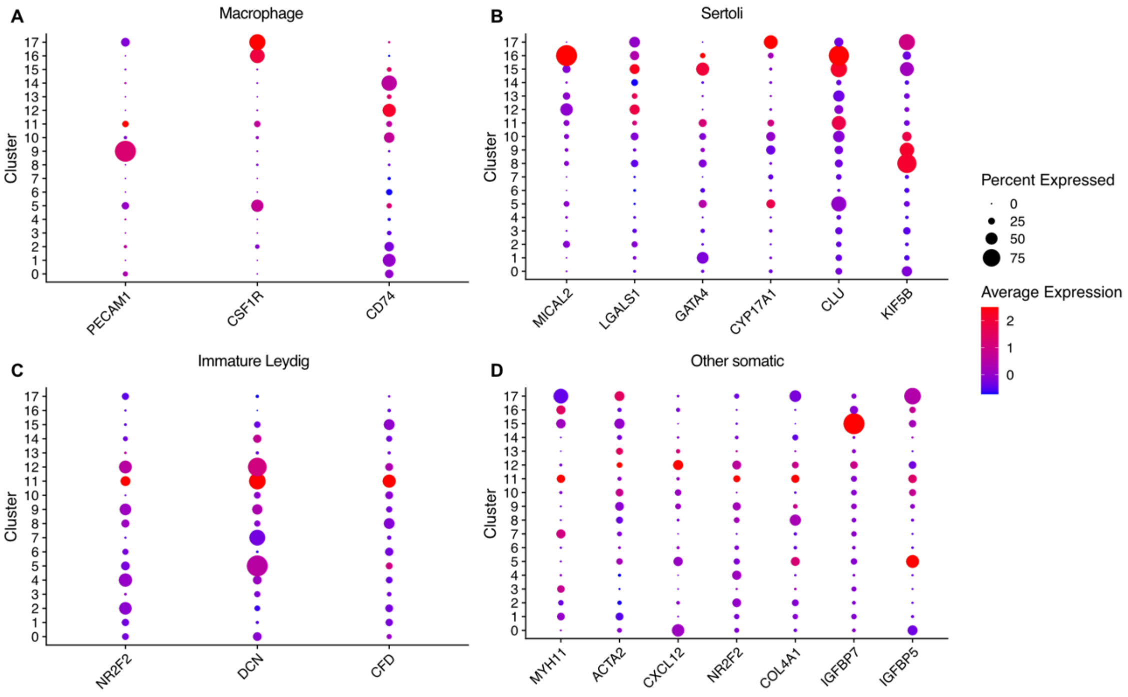


**Supplementary Figure 2. Average gene expression of mammalian testis marker genes and percent of cells where each gene was expressed per testis cell cluster in flycatchers.** The Y-axis show known mammalian gene markers for (A) macrophages, (B) Sertoli cells, (C) Immature Leydig cells and (D) other somatic cells.

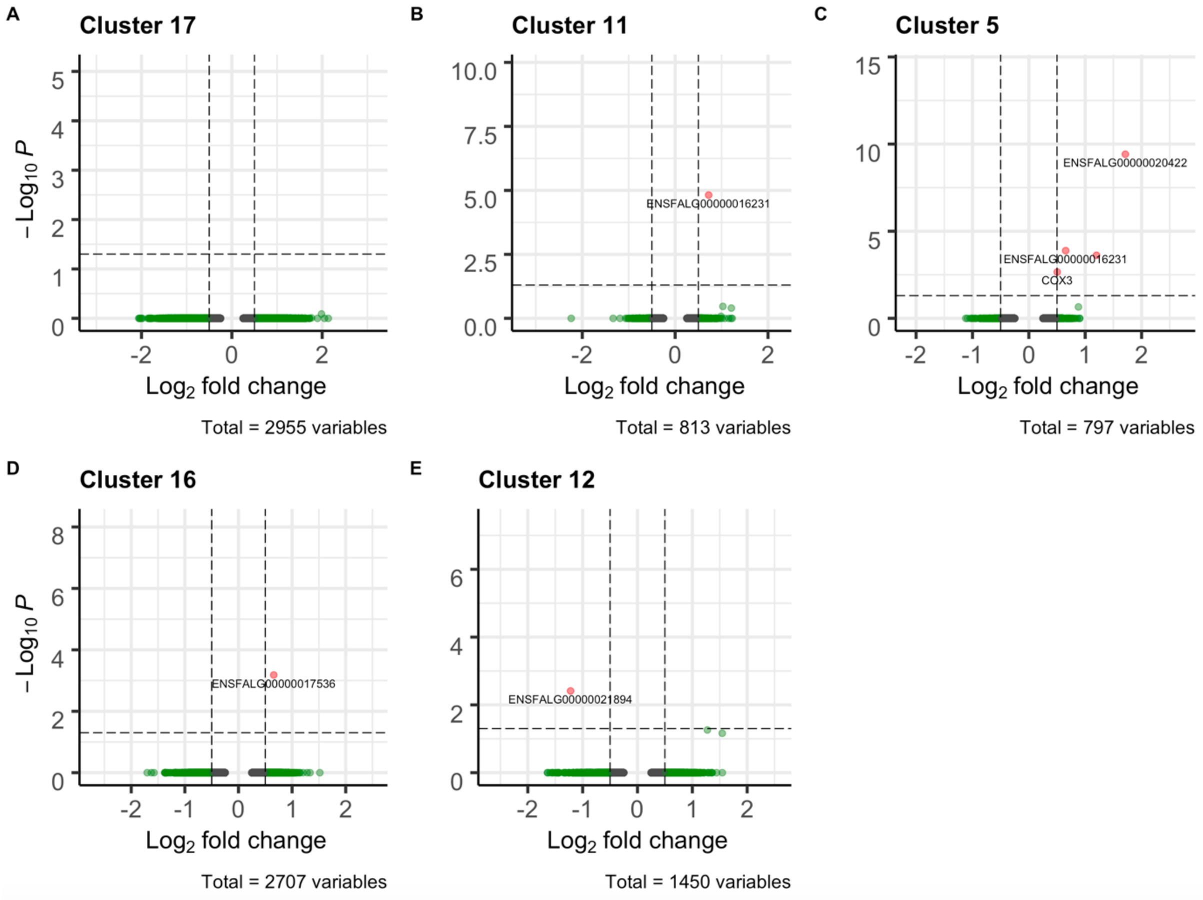


**Supplementary Figure 3. Volcano plots showing differences in fold change between collared and pied flycatchers for somatic cell clusters obtained from single-cell RNA sequencing of testis.**

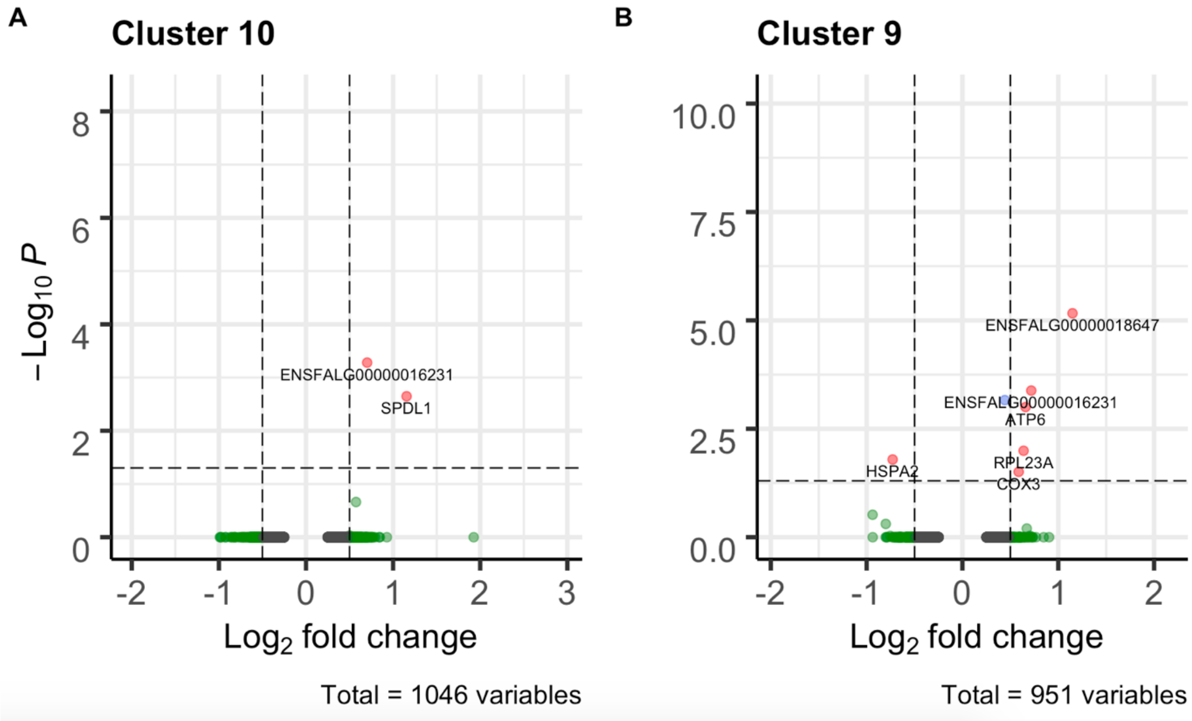


**Supplementary Figure 4. Volcano plots showing differences in fold change between collared and pied flycatchers for spermatogonia cell clusters obtained from single-cell RNA sequencing of testis.**

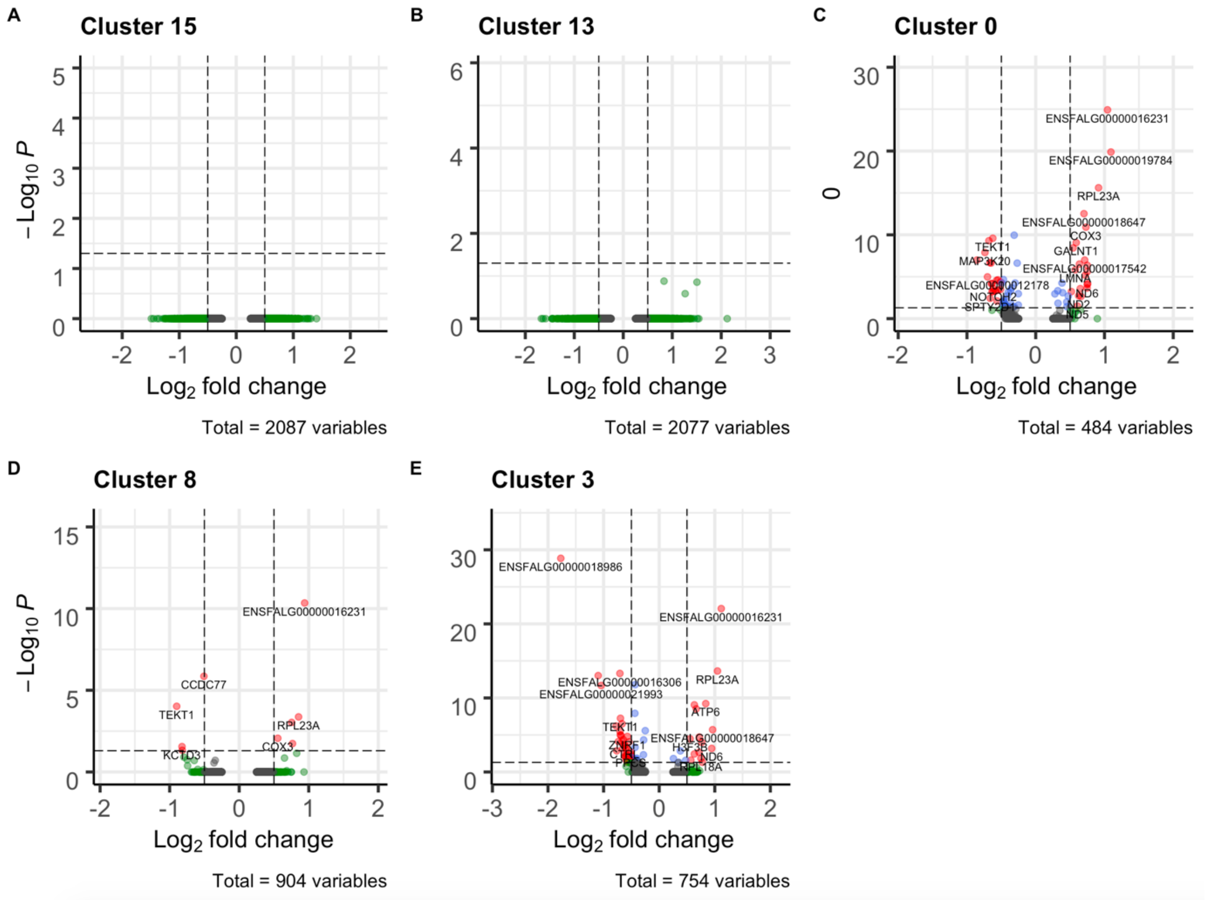


**Supplementary Figure 5. Volcano plots showing differences in fold change between collared and pied flycatchers for spermatocyte cell clusters obtained from single-cell RNA sequencing of testis.**

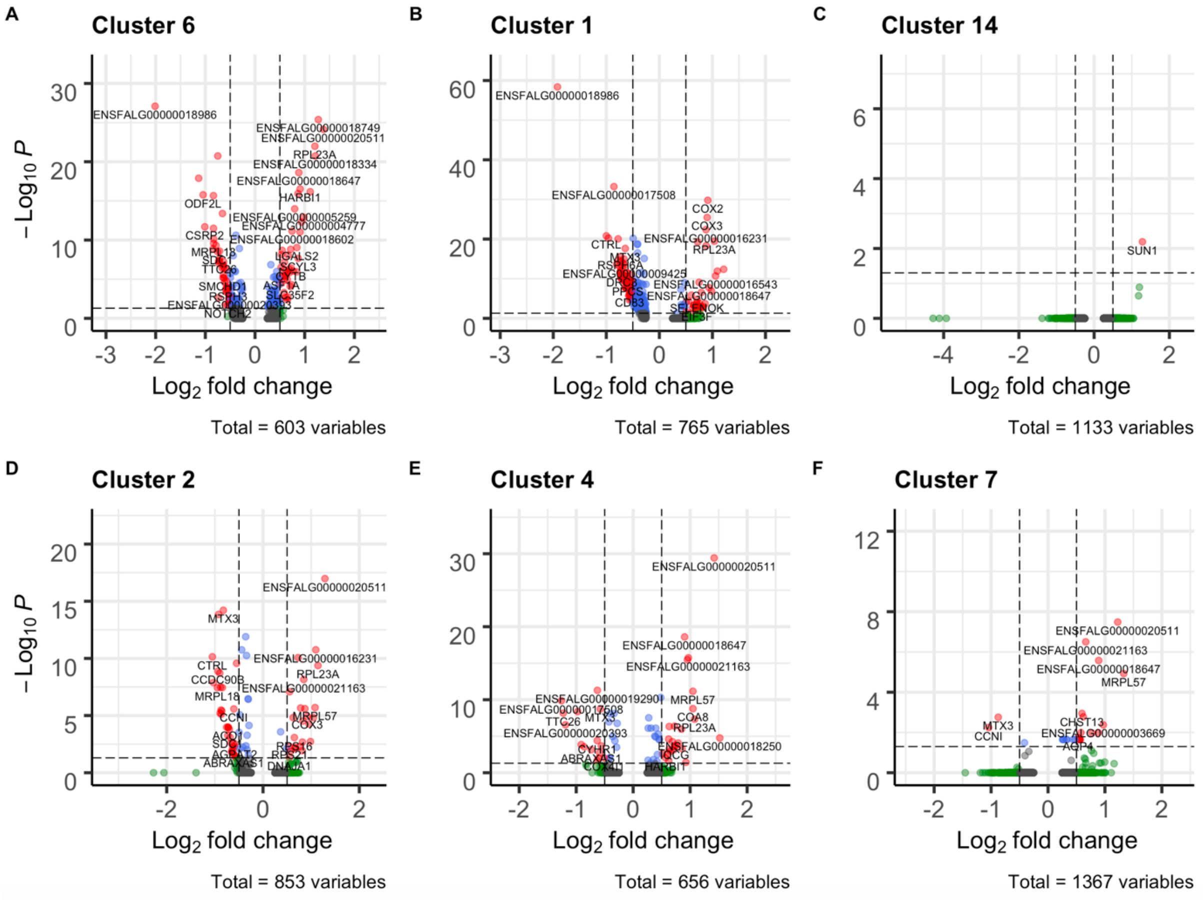


**Supplementary Figure 6. Volcano plots showing differences in fold change between collared and pied flycatchers for spermatid cell clusters obtained from single-cell RNA sequencing of testis.**

Supplementary table 1. Gene Markers for all clusters. (Separate file).

Supplementary table 2. Chi square contingency table for private and shared markers of *Ficedula* flycatchers by stage. ($\chi^2$ = 76.411, df=2, $p<2.2e^{-16}$)

| | Number of gene markers | |
|---|---|---|
| Stages | Shared | Private |
| Spermatogonia | 593 | 965 |
| Spermatocytes | 278 | 755 |
| Spermatids | 46 | 252 |

Supplementary table 3. Post hoc analysis for Pearson's Chi squared results. Significant p values are in bold.

| Dimension | Value | Shared | Private |
|---|---|---|---|
| Spermatogonia | Residuals | 7.90 | -7.90 |
| Spermatogonia | P values | **0** | **0** |
| Spermatocytes | Residuals | -4.16 | 4.16 |
| Spermatocytes | P values | **1.9^10$^{-4}$** | **1.9^10$^{-4}$** |
| Spermatids | Residuals | -6.39 | 6.39 |
| Spermatids | P values | **0** | **0** |

Supplementary table 4. Average expression of genes per cluster. (Separate file)

Supplementary table 5. Enrichment analysis of genes expressed in the Z chromosome versus the autosomes per cluster for all expressed genes > 0.01

| Stage | Cluster | Autosomes | | Z Chromosome | | P-value |
|---|---|---|---|---|---|---|
| | | Expressed | Non-Expressed | Expressed | Non-Expressed | |
| Somatic cells | 17 | 5591 | 8039 | 247 | 343 | 0.3566 |
| Somatic cells | 11 | 8810 | 4820 | 412 | 178 | **0.0051** |
| Somatic cells | 5 | 9151 | 4479 | 422 | 168 | **0.014** |
| Somatic cells | 16 | 7472 | 6158 | 352 | 238 | **0.0114** |
| Somatic cells | 12 | 6996 | 6634 | 325 | 265 | **0.0403** |
| Spermatogonia | 10 | 8182 | 5448 | 386 | 204 | **0.0047** |
| Spermatogonia | 9 | 8334 | 5296 | 384 | 206 | **0.0294** |
| Meiosis stages | 15 | 7512 | 6118 | 350 | 240 | **0.0241** |
| Meiosis stages | 13 | 7212 | 6418 | 327 | 263 | 0.1241 |
| Meiosis stages | 0 | 8580 | 5050 | 387 | 203 | 0.1036 |
| Meiosis stages | 8 | 8296 | 5334 | 369 | 221 | 0.2198 |
| Meiosis stages | 3 | 8610 | 5020 | 386 | 204 | 0.1426 |
| Spermatids | 6 | 7683 | 5947 | 343 | 247 | 0.2105 |
| Spermatids | 1 | 8431 | 5199 | 379 | 211 | 0.1305 |
| Spermatids | 14 | 6417 | 7213 | 298 | 292 | 0.0558 |
| Spermatids | 2 | 7788 | 5842 | 358 | 232 | **0.0482** |
| Spermatids | 4 | 7372 | 6258 | 324 | 266 | **0.0045** |
| Spermatids | 7 | 7622 | 6008 | 348 | 242 | 0.0768 |

Supplementary table 6. Enrichment analysis of Z-linked genes versus autosome genes per cell cluster for the top 500 expressed genes.

| Stage | Cluster | Autosomes | | Z Chromosome | | P-value |
|---|---|---|---|---|---|---|
| | | Expressed | Non-Expressed | Expressed | Non-Expressed | |
| Somatic cells | 17 | 473 | 13157 | 27 | 563 | 0.0977 |
| Somatic cells | 11 | 475 | 13155 | 25 | 565 | 0.1931 |
| Somatic cells | 5 | 470 | 13160 | 30 | 560 | **0.0275** |
| Somatic cells | 16 | 479 | 13151 | 21 | 569 | 0.5093 |
| Somatic cells | 12 | 476 | 13154 | 24 | 566 | 0.2583 |
| Spermatogonia | 10 | 470 | 13160 | 30 | 560 | **0.0275** |
| Spermatogonia | 9 | 466 | 13164 | 34 | 556 | **0.0033** |
| Meiosis stages | 15 | 475 | 13155 | 25 | 565 | 0.1931 |
| Meiosis stages | 13 | 477 | 13153 | 23 | 567 | 0.3345 |
| Meiosis stages | 0 | 476 | 13154 | 24 | 566 | 0.2583 |
| Meiosis stages | 8 | 473 | 13157 | 27 | 563 | 0.0977 |
| Meiosis stages | 3 | 480 | 13150 | 20 | 570 | 0.6003 |
| Spermatids | 6 | 480 | 13150 | 20 | 570 | 0.6003 |
| Spermatids | 1 | 478 | 13152 | 22 | 568 | 0.4193 |
| Spermatids | 14 | 474 | 13156 | 26 | 564 | 0.1397 |
| Spermatids | 2 | 479 | 13151 | 21 | 569 | 0.5093 |
| Spermatids | 4 | 480 | 13150 | 20 | 570 | 0.6003 |
| Spermatids | 7 | 480 | 13150 | 20 | 570 | 0.6003 |

Supplementary table 7. Differential expression analysis between Collared and Pied flycatcher per cell cluster. (Separate file).

Supplementary table 8. General linear model with binomial distribution having as response variable DE genes vs non-DE genes per cluster and spermatogenesis stage (ie: somatic cells, mitosis, meiosis or spermiogenesis) as an explanatory variable. Number of observations: 18.

| | Estimate | S.E. | Z value | P value |
|---|---|---|---|---|
| Intercept | -4.21277 | 0.07426 | -56.726 | **< 0.001** |
| Somatic cells | -17.79283 | 275.73098 | -0.065 | 0.949 |
| Spermiogenesis | 1.81224 | 0.08207 | 22.082 | **< 0.001** |
| Mitosis | -2.28651 | 0.41525 | -5.506 | **< 0.001** |

Supplementary table 9. Significant GO terms of DE genes between collared and pied flycatchers. (Separate file).
